# Human lncRNAs harbor conserved modules embedded in different sequence contexts

**DOI:** 10.1101/2023.11.01.565086

**Authors:** Francesco Ballesio, Gerardo Pepe, Gabriele Ausiello, Andrea Novelletto, Manuela Helmer-Citterich, Pier Federico Gherardini

**Affiliations:** PhD Program in Cellular and Molecular Biology, Department of Biology, University of Rome “Tor Vergata”, Rome, Italy; Department of Biology, University of Rome “Tor Vergata”, Rome, Italy

## Abstract

We analyzed the structure of human long non-coding RNA (lncRNAs) genes to investigate whether the non-coding transcriptome is organized in modular domains, as is the case for protein-coding genes. To this aim, we compared all known human lncRNA exons and identified 340 pairs of exons with high sequence and/or secondary structure similarity but embedded in a dissimilar sequence context. We grouped these pairs in 106 clusters based on their reciprocal similarities. These shared modules are highly conserved between humans and the four great ape species, display evidence of purifying selection and likely arose as a result of recent segmental duplications. Our analysis contributes to the understanding of the mechanisms driving the evolution of the non-coding genome and suggests additional strategies towards deciphering the functional complexity of this class of molecules.

**Author summary:** The Human genome includes more than 18,000 genes coding for RNAs that are not translated into proteins, called long non-coding RNAs (lncRNA). Mounting evolutionary and experimental evidence shows that a large amount of these RNAs have a specific function, mainly as regulators of a diverse set of biological processes. Here we set out to investigate whether these genes have a modular organization similar to that of protein-coding genes. Accordingly, we compared the sequence of all the exonic regions of human lncRNAs and identified 106 clusters of non-repetitive exonic modules shared between this class of genes. These modules display evidence of purifying selection, are highly conserved between humans and the four great ape species, and may represent distinct functional units that have been shuffled among multiple lncRNA genes, in a manner similar to the exon-shuffling process that is observed in the coding genome.

## Introduction

Many eukaryotic proteins are composed of a discrete number of domains, endowed with autonomous folding capacity and/or characteristic functions. This type of organization is defined as modular, and the process by which this set of modules is recombined into a variety of different protein products is known as “exon-shuffling” [1].

Long noncoding RNAs (lncRNAs) represent a heterogeneous class of RNAs that are not translated into functional protein products but, similar to messenger RNAs, are transcribed from genes that may have an exon/intron structure. These RNAs are generally defined as non-coding RNAs of more than 200 nucleotides in length and can be capped, polyadenylated and spliced [2], much in the same way as the transcripts of protein coding genes. The human genome contains about 18,000 lncRNA genes and 47,000 transcripts [3], most of which are of unknown function. lncRNAs exhibit evidence of purifying selection and experimental evidence shows that at least a portion of them is indeed functional (287 eukaryotic lncRNAs associated with a biological function are collected by [4], 1,273 human lncRNAs by [5]). Some lncRNAs have been characterized in depth and they may function as regulatory molecules both in the nucleus and the cytoplasm, through a variety of mechanisms, including interaction with transcription factors, recruitment of chromatin modifying complexes, modulation of the expression of their neighboring genes, control of mRNA stability and translation and competition for the binding of specific miRNAs [6–8]. Individual lncRNAs have been found to have a role in promotion of metastasis [9], neuronal differentiation [10], regulation of the accumulation of beta amyloid peptide in Alzheimer’s disease [11], and many other processes in a diverse array of pathological and physiological contexts. However the identification of the function of lncRNAs on a global scale remains elusive [12], also because their definition likely encompasses an extremely heterogeneous set of genes, whose main, and possibly only, common characteristic is the fact that they do not produce a functional protein product [13].

In general, lncRNAs are significantly less conserved than protein-coding sequences [14], which also suggests that the relationship between sequence and function is particularly complex in this class of molecules. Examples of lncRNA such as *Xist, Megamind, Cyrano* and *Miat* have been described, which have conserved functions throughout multiple organisms, and yet display a level of sequence divergence that challenges sequence homology search tools [13,15]. A corollary of this observation is that similarity amongst lncRNA within a given organism is also limited, and, unlike coding sequences, most lncRNAs appear in single copies in vertebrate genomes [13].

However, lncRNAs are significantly more likely to contain repetitive sequences, particularly transposable elements (TEs) [15,16]. On one hand, this could simply indicate that lncRNAs are more prone to transposon insertion, because of their aforementioned looser association between sequence and function [13]. On the other hand, this observation implies the existence of stretches of homologous sequences that are shared among different lncRNAs, even when the lncRNAs themselves are not related by descent.

Because TEs are often enriched in sequences with regulatory function, and may contribute to their “spread” within a genome [17], Johnson and Guigò [18] hypothesized that the presence of TEs may result in the sharing of functional cassettes among evolutionarily unrelated lncRNA, possibly implying a modularization of function for this class of molecules [6,12], reminiscent of the notion of domains in the protein-coding world. In support of this hypothesis, it has been reported that TE-derived sequences within lncRNAs are more conserved compared with non-TE sequences [19].

Here we set out to expand the identification of modules in lncRNAs that could have contributed to increasing the diversity of the non-coding genome, similar to the exon-shuffling phenomenon that is well known for protein sequences. Our work extends previous observations in three ways, namely by i) focusing on the sharing of individual exons among unrelated lncRNAs within the human genome, ii) specifically excluding exons that contain repetitive sequences, and iii) including secondary structure as an additional criterion to define similarity, as lncRNAs with similar functions often lack linear sequence homology [20], and many examples of ncRNA are known whose function is tied to their secondary structure [21–24].

## Results

### Exon sequence and secondary structure comparison

In order to search for similarities among lncRNAs, we performed a pairwise comparison of both the sequence and the predicted secondary structure of 12,097 non-overlapping human lncRNA exons that do not contain repetitive sequences, performing a total of more than 73 million sequence alignments and an equal number of structure alignments. The distributions of the corresponding scores are shown in Figure 1A, B (1A-B Fig.).

**Fig.1.**
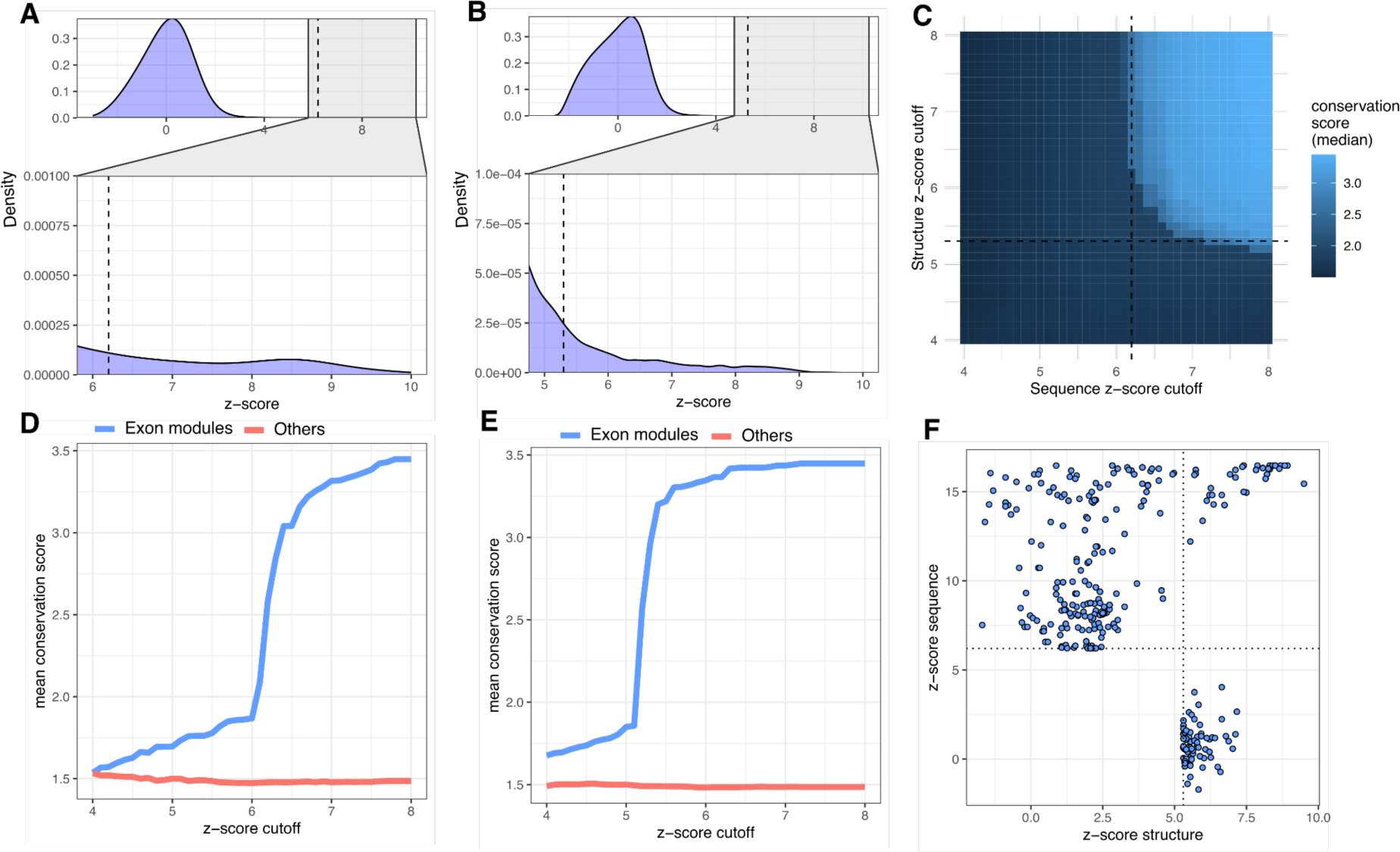
Sequence and structure alignments results. A) Distribution of z-transformed pairwise alignment scores for sequence; B) Distribution of z-transformed pairwise alignment scores for structures, for these distributions, a close-up around the proposed cutoff thresholds is also shown; C) heatmap representing the conservation scores in the four non-human primates of all pairs selected at the different z-score thresholds of sequence and structure alignments; D, E) Mean conservation scores (within four non-human primates) of members of clusters defined by different z-score thresholds of pairwise similarity for sequence (D) and structure (E). Note the steep increase in evolutionary conservation for the z-score cutoff of 6.2 (sequence) and 5.3 (structure), respectively; F) Scatter plot of sequence and structure similarity z-scores of the exon pairs (for the sake of clarity, the more than 73 million pairs below the thresholds are not shown).

To identify pairs or groups of exons representing shared sequence elements, hereafter referred to as “modules”, it was necessary to select a threshold above which their sequence or structure similarity would be considered significant.

We thus investigated the conservation of lncRNA exons in four non-human primates (see Materials and Methods), with the goal of identifying shared sequence elements in the human genome that are also conserved in other primate genomes.

Accordingly, we calculated the mean conservation scores of sequence modules across these species, as a function of the similarity score threshold used to define the modules themselves. Using this procedure, we observed a sharp transition in conservation at Z-score similarity thresholds of 6.2 and 5.3 for sequence and structure alignments, respectively (1C-D-E Fig.). We consider this increase in conservation, coupled with the high Z-score similarity threshold, as a strong indication that the shared sequence elements we identified represent significant similarities. As a further benchmark, we repeated the entire procedure by aligning exons against random sequences with the same length and base composition. None of the alignments produced z-scores above the 6.2 threshold.

By using these thresholds, we identified a total of 340 exon pairs (219 identified by sequence, 75 by structure and 46 by both), involving 338 different exons and 218 different genes (1F Fig.). Starting from these pairwise similarities, we identified 106 clusters (exon modules) defined by homologous lncRNA exons represented in at least two copies in the same or different genes (2 Fig., S1 Fig. and S1 Table).

**Fig.2.**
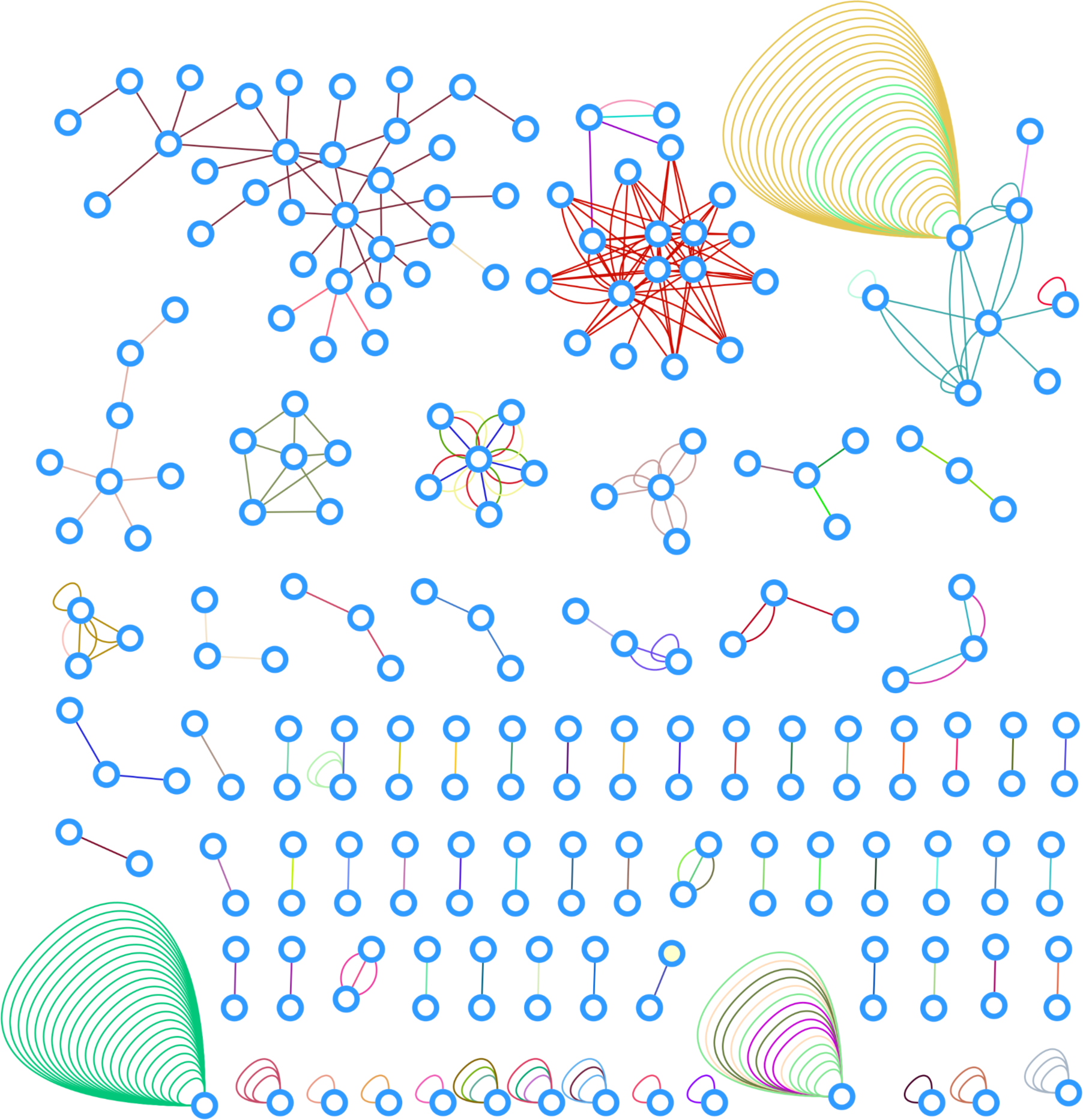
Network representation of the exon-sharing gene clusters and the corresponding exon modules. Each node represents a lncRNA gene and each edge an exonic module shared between two genes. Same color edges within a gene cluster represent a module. Self-loops represent instances where the same module occurs multiple times in a single gene. The network representation was generated using Cytoscape [25].

To rule out the possibility that similarity between exons in a pair of genes is simply due to paralogy, we aligned the entire genes using BLAST and excluded pairs with alignment coverage on the smallest gene of the pair greater than 80%. Measuring the alignment coverage of the entire genes, including introns, allowed us to identify and exclude cases of complete paralogy even in the presence of intronization or imprecise exon annotation.

We note that, in general, our analysis is dependent on the reliability of the reconstruction of the whole transcript structure, which is used to define the exons themselves. This is summarized by the Transcript Support Level (TSL, S1 Table).

Figure 3A (3A Fig.) is an example of one of the identified exon modules shared by a group of 7 lncRNA genes: ENSG00000279072.1, ENSG00000188185.11, ENSG00000276997.4, ENSG00000280136.2, ENSG00000280279.1, ENSG00000230724.9, ENSG00000238035.8. This cluster consists of 9 exons that contain a region of ∼65 nucleotides with high sequence similarity (external gap trimmed sequence identity 92-98%, 3B Fig.) embedded in different genes. It is worth noting that, in some cases, the module constitutes an exon on its own, whereas in other cases it is part of a larger exon.

**Fig.3.**
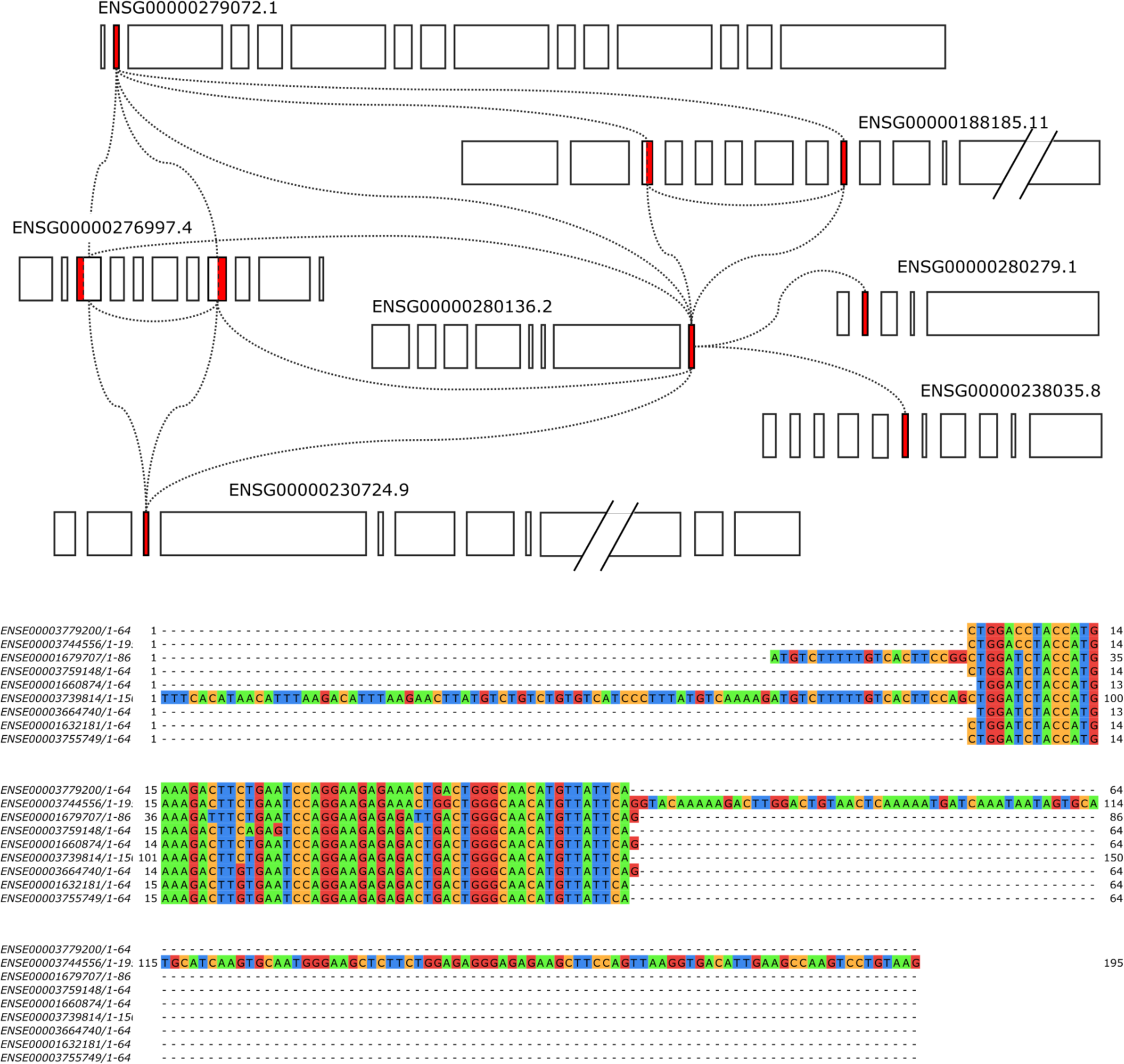
An example of the identified exon modules. A) Schematic representation of 7 genes containing representatives (in red) of exons contributing to a module cluster. Each box represents an exon, with width proportional to its length (intron length not to scale); B) multiple alignment of the 9 exons contributing to the cluster.

We then analyzed in more detail the sequence context of exon modules. More specifically, we looked at the sequence similarity of additional exons flanking the modules, to rule out the possibility that the similarity between modules in different genes simply reflects global sequence similarity between the exonic components of genes (see Materials and Methods and 4A Fig.). The alignment scores for exons flanking the putative module in the same gene, upstream and downstream (4B Fig.), showed that the similarity between exon modules is significantly higher than that of the sequence context in which they are embedded. We also observed a small proportion of cases in which the flanking exons are also similar (outliers in 4B Fig.). These cases fall outside the criteria used to define exon modules: in 17 cases because they are less than 50 nucleotides in length, and in another 17 cases because they contain repetitive sequences.

**Fig.4.**
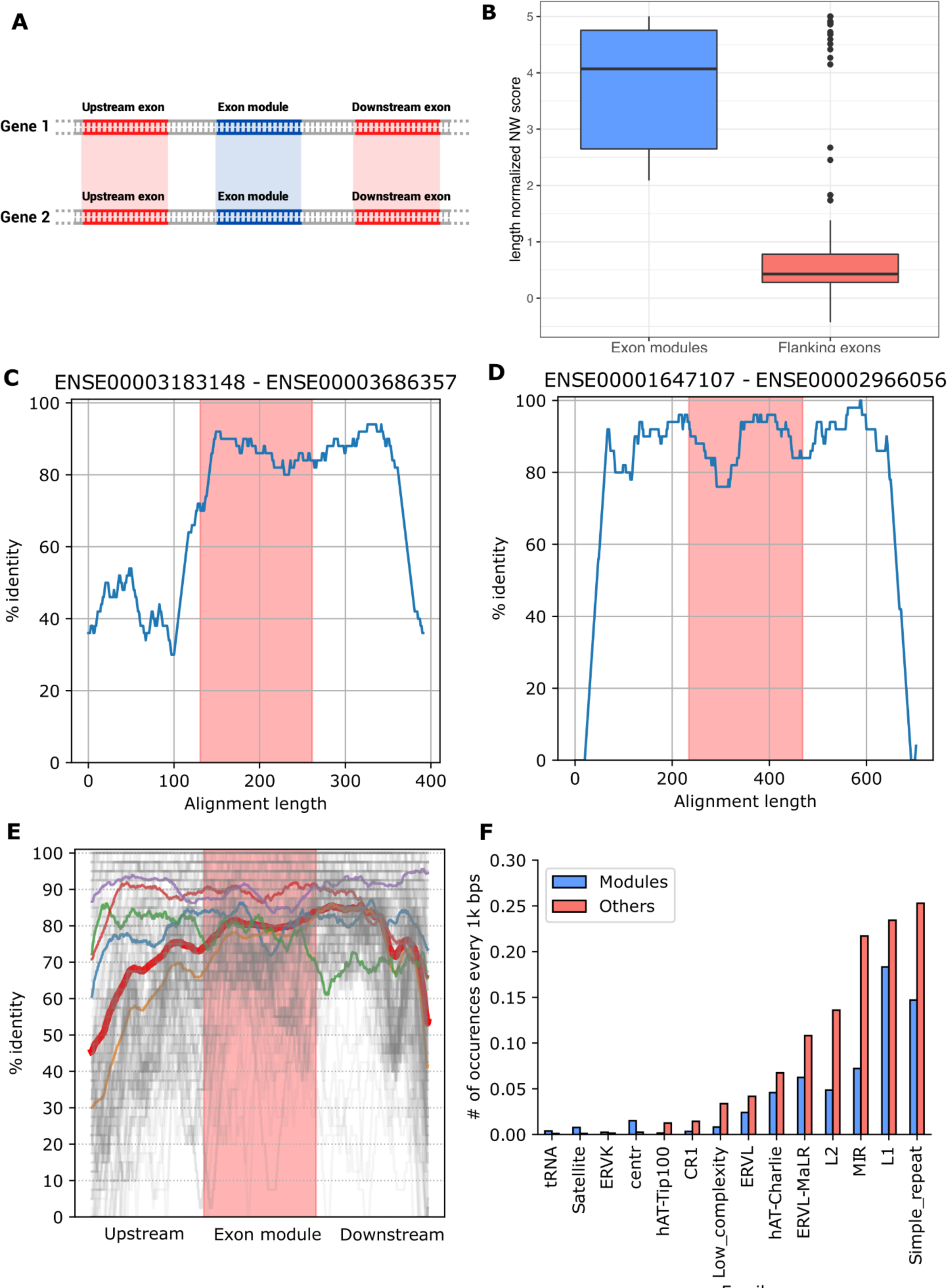
Analysis of the sequence regions flanking exon modules. A) For each pair of genes containing a shared exon module we compared the similarities of the upstream and downstream flanking exons (when present); B) Distributions of the length-normalized Needleman and Wunsch scores of exonic modules (in blue) and of their upstream and downstream flanking exons (in red); C) A pair of exons in which the similarity only extends to the downstream flanking intron; D) A pair of exons in which the similarity extends upstream and downstream into both flanking introns; E) Overall representation of all the length-scaled similarities between all the exon pairs and their flanking introns (in grey), the median identity percentage is represented in red. The other colored lines represent five clusters of similarity patterns as defined by grouping individual lines; F) Number of occurrences per thousand base pairs of families of repetitive sequences in flanking introns with significant differences (padj<0.05) between the exonic modules and the other lncRNA exons.

We then analyzed the sequence similarity of the intronic sequences flanking the exon modules. To this end, we defined genomic regions of interest by extending upstream and downstream the sequence of each candidate exon pair, until we obtained two sequences with a length equal to three times that of the longest exon of the pair (4C-E Fig.). We limited the analysis to pairs with sequence similarity above the z-score threshold of 6.2 and excluded modules repeating within the same gene. For each pair of genomic regions of interest, we performed a global alignment using the same parameters used to identify the exon modules, and calculated the percentage identity of the pairs using overlapping windows of 50 nucleotides with a single nucleotide shift, to generate graphs depicting the extent of the similarity. We found that, in the majority of instances, sequence similarity extends into the flanking intronic regions. More specifically, in approximately one third of the cases, the similarity encompassed both the upstream and downstream intron, in another third of the cases the similarity extended to a single intron, while the remainder of cases lacked a clear pattern. We did not observe any cases where the similarity was confined to the boundaries of the candidate exon modules.

The extension of the similarity through the flanking introns suggests that the most common mechanism responsible for the origin of exon modules is segmental duplication of a genomic DNA stretch encompassing the parental copy of an exon. This is the same mechanism suggested as a driver of exon shuffling in protein coding genes [26]. To further confirm these findings, we compared our results with the data present in the UCSC Segmental Dups track (genomicSuperDups) which contains regions detected as putative genomic duplications within the human genome. These regions represent large recent duplications (>= 1 kb and >= 90% identity) that originated over the last ∼40 million years of human evolution, based on neutral expectation of divergence [26]. For 84 of the 340 lncRNA exon pairs identified here, we found a match in the segmental duplications identified by Bailey et al. In 81 of these cases the duplicated stretch includes the entire exons of the pair, while in 3 cases the duplication is interrupted within the exon. We also observed a higher frequency of pairs located on the same chromosome (∼20.5%) compared with what is observed when the same exons are randomly paired (∼3.6%). Moreover, pairs of exon modules that are on the same chromosome are closer together when compared to the same random pairing control (Mann-Whitney p-value=9.86e-05). A higher rate of occurrence on the same chromosome has been described for segmental duplications [27]. To further extend the analysis of flanking regions, we compared the rate of occurrence of multiple families of repetitive elements in the introns flanking candidate exonic modules vs other lncRNA exons (for exons located at the ends of a gene, we included a region of 10k bps in the genome). We calculated the number of occurrences per 1,000 base pairs of each family of repetitive elements on the set of regions flanking the exon modules vs the other lncRNA exons (4F Fig., S2 Table) thus obtaining a distribution of occurrences where the observations correspond to the individual sequence regions. We then compared these distributions using a Mann-Whitney U test, with Bonferroni correction for multiple hypothesis testing. We observed significant differences for 15 of 46 families (padj<0.05). Interestingly, centromere and satellite repeats are among the few classes of repeats enriched in regions flanking the exon modules, while most classes of transposon-or endogenous retrovirus-derived repeats are depleted. Since the genomic regions proximal to centromeres and telomeres are enriched with segmental duplications [28], this observation further points at segmental duplication as the main driver of the appearance of these exon modules, as opposed to, for instance, transposition. The enrichment of this type of repetitive sequences can be explained by the localization near the centromeres or telomeres of a portion of the modules (S2 Fig.). Moreover, searching for transposase domains using a procedure similar to the one described in [29] did not reveal significant differences in their occurrence among genes containing exon modules (data not shown), further highlighting that transposition is not the main driver of this process.

To further investigate the characteristics of these modules we looked at the distribution of cis-regulatory elements (CRE) within their sequences (5A Fig.). This research highlighted a depletion in exon modules of the most frequent CREs (Fisher’s exact test padj=6.2e10-3, 1.2e10-2, 1.2e10-2 for pELS, dELS and PLS, CTCF-bound respectively). One of the few elements that are not depleted are H3K4me3 marks, which are characteristic of transcriptionally active regions (Howe et al. 2017). Interestingly this histone modification is usually found in the region corresponding to the beginning of the transcript [30]. Accordingly, when we investigated the position of the exonic modules within their transcripts (5B Fig.), we detected a higher frequency of the modules at the 5’ end. This finding is consistent with what is observed in protein coding genes, which in vertebrates tend to increase their length over time by gaining recently evolved domains, primarily through the addition of sequences at the 5’ end of genes [31]. The insertion of these modules at the extremities of the transcript presumably allows the addition of genetic material with minimal disruption to the existing sequence.

**Fig 5.**
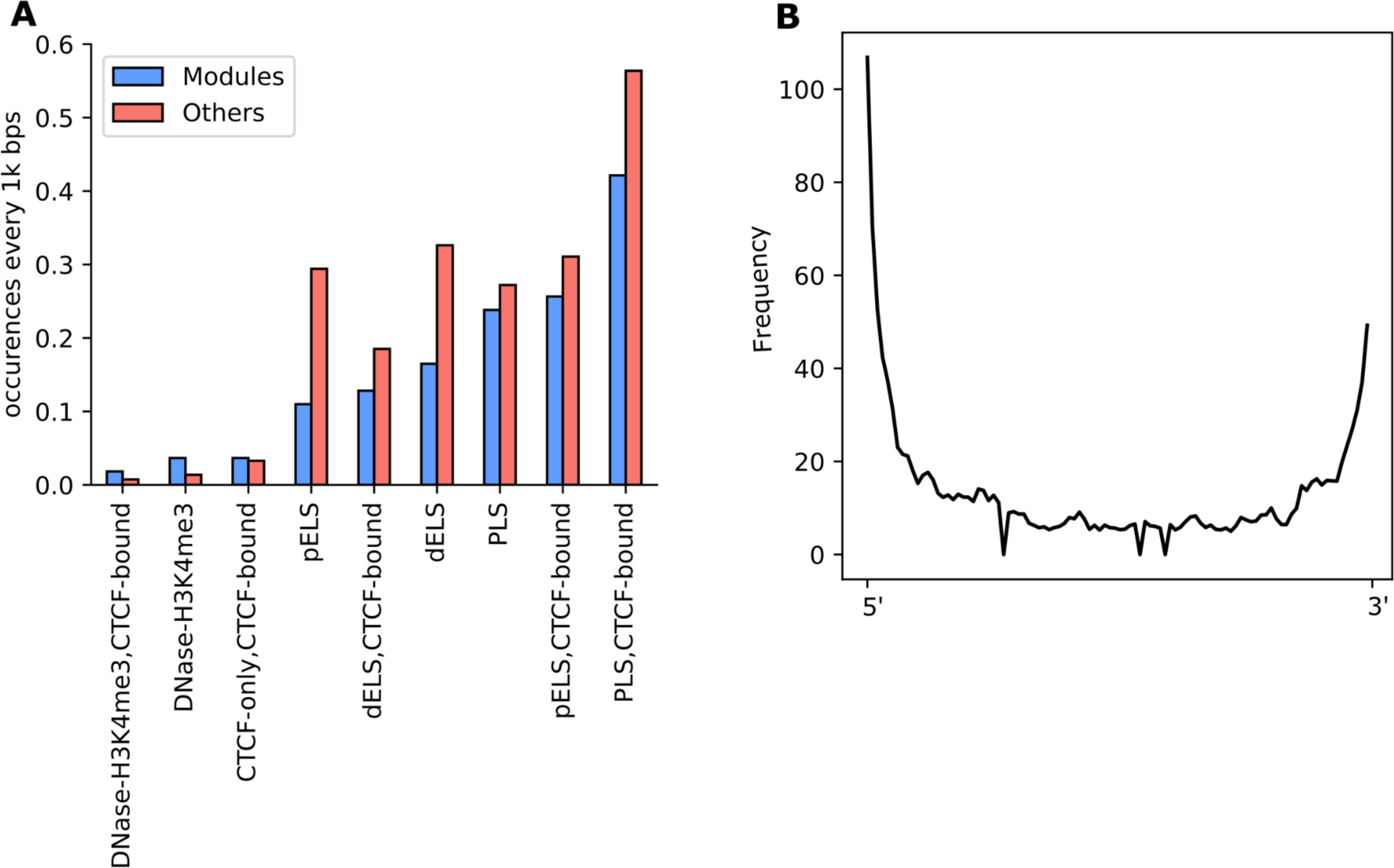
Cis-regulatory elements (CRE) and position of the modules. A) number of occurrences of the different CREs from the annotation present in ENCODE every thousand nucleotides in the modules (in blue) and in the other lncRNA exons of the dataset (in red); B) the y axis indicates the frequency of regions containing modules relative to their position on their transcript (which is indicated on the Y axis, see Methods), as the sum of modules present in that region. The higher y value therefore indicates that there is a greater number of modules at the ends of the transcripts, particularly at the level of the 5’ end.

### Evolutionary conservation of exon modules

To analyze in detail the inter-specific conservation of exon modules, we compared their conservation scores (see Materials and Methods) with the conservation scores of functionally annotated lncRNA exons, using the conservation scores of other lncRNA exons as control. Functionally annotated lncRNA genes were collected from the lnc2Cancer database [5], which contains experimentally supported annotations of lncRNA associated with a biological function, as derived from the literature (see Materials and Methods). The comparison of these three categories revealed that the conservation score of exon modules was higher than that of exons belonging to functionally annotated lncRNA genes (Mann-Whitney p-value=6.3e-5), and both the conservation score of exon modules and of exons belonging to functionally annotated lncRNA were significantly higher than the conservation score of the remaining lncRNA exons (Mann-Whitney p-value=7.4e-27 and 3.5e-26 respectively, 6A Fig.). When looking at the conservation of exon modules in four higher primate species, we also observed a greater proportion of exons with a BLAST hit among exon modules vs the remaining exons. More specifically, 65.97% of the exon modules have a BLAST hit in Chimpanzee, 42.01% in Bonobo, 11.83% in Gorilla and 47.04% in Orangutan. Conversely, only 43.23%, 16.72%, 2.61%, 25.86% of the control exons (i.e the portion of the 12097 lncRNA exons that have no repetitive and non-overlapping sequences and that are not modules) have BLAST hits on the same species respectively (6B Fig.) To evaluate the significance of these results we performed a Fisher’s exact test on the aggregated data from the different species, which confirmed that these results are significant (p-value=4.10e-15).

**Fig.6.**
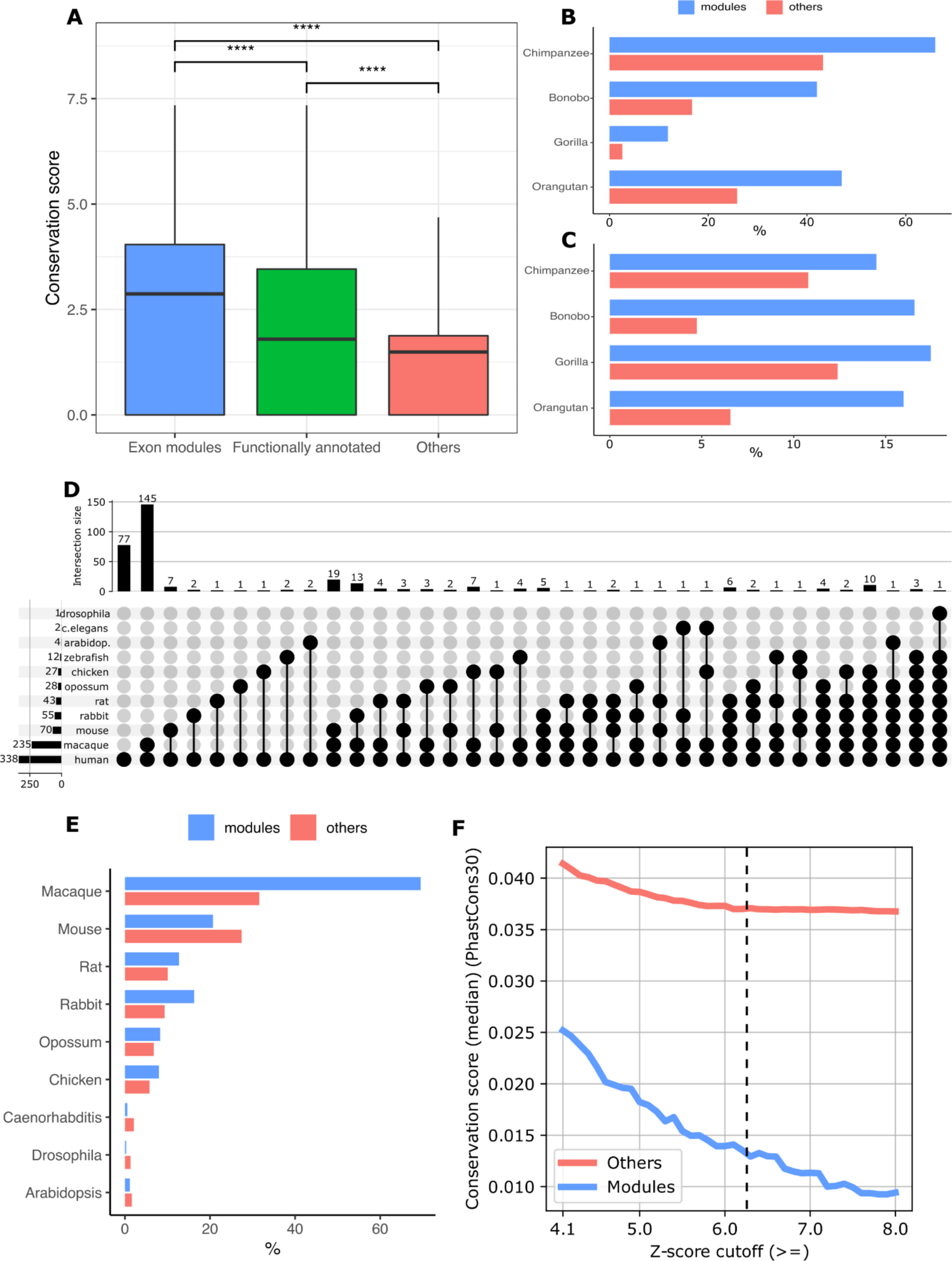
Evolutionary conservation of exon modules. A) Box-plot of the conservation scores in four non-human primates for exon modules, functionally annotated exons from the lnc2Cancer database, and controls; B) Percentage of exon modules (in blue) and other exons (in red) that showed a BLAST hit (e-value <0.001) in the primate species considered; C) Percentage of genes showing a conserved syntenic region (as defined in SynthDB) among those containing exon modules (in blue) vs genes not containing an exon module (in red); D) Upset plot representing the exons that have a BLAST hit in the species analyzed in Sarropoulos et al. and in other model organisms; E) Percentages of modules (in blue) and other exons (in red) showing a BLAST hit in the indicated species F) PhastCons 30 mammals scores of members of clusters defined by different z-score thresholds of pairwise similarity from sequence alignments (in blue) and the other lncRNA exons of the dataset (in red).

Since the BLAST similarity score with non-human primates does not take into account the genomic position of exons in different organisms, i.e. it cannot distinguish between the similarity of true orthologs vs in- and out-paralogs, we investigated whether exon modules are located in regions of synteny between non-human primates more often than other exons. To this end we leveraged the SynthDB [32] database, which provides data on orthology relationships between humans and other primates. We observed that the percentage of genes located in a syntenic region is higher for genes that contain at least one exon module, compared with those which do not. Accordingly, 14.50% of the exon modules are located in genes that have an ortholog in Chimpanzee, 16.57% in Bonobo, 17.46% in Gorilla and 15.98% in Orangutan. While for the other lncRNA exons we observed percentages of 10.79, 4.74, 12.39, 6.55 in the same species respectively. We then performed a Fisher’s exact test comparing exons modules that belong to genes with an ortholog in at least one of the species mentioned above to the other exons which confirmed the significance of our results (p-value=5.63e-05) (6C Fig.).

To strengthen the evolutionary conservation analysis, and to compare our results with the analysis by Sarropoulos et al. 2019 [33], we extended it by including additional species. To this end, we aligned all lncRNA exons using blastn against the genomes of the organisms used in Sarropoulos et al. 2019 (Macaque, taxid: 9544; Rabbit, taxid: 9986; Chicken, taxid: 9031; Opossum, taxid: 13616; Rat, taxid: 10116; Mouse, taxid: 10090), and other model organisms (Danio rerio, taxid: 7955; Drosophila melanogaster, taxid: 7227; Caenorhabditis elegans, taxid: 6239; Arabidopsis thaliana, taxid: 3702), using an e-value threshold of 0.01 to identify hits (5D-E Fig., S3 Fig.). Figure 6E (6E Fig.) displays the percentage of exonic modules vs other lncRNA exons that have at least one hit in the species indicated above. This analysis shows a rapid decay in the number of similar exons as the evolutionary distance from humans increases. Figure 5F (5F Fig.) shows the 30 mammal PhastCons scores of the exon modules, as a function of the z-score similarity threshold used to define the modules themselves (i.e. the threshold described in 1A Fig.). This analysis demonstrates that the exon modules identified in this work, which are highly similar as they were selected on the basis of having a Z-score of at least 6.2 and 5.3 in the sequence and structure alignment respectively, represent duplications that are recent (as implied by the high levels of sequence similarity) and that are exclusively found in humans and higher primates, and thus have lower PhastCons scores on the entire set of 30 mammals (6F Fig.).

Overall, the above results reveal that roughly 4% of lncRNA genes (218 lncRNA genes/5,423 total lncRNAs genes which contain at least one exon without repetitive sequences, see Materials and Methods) include one or more exons having significant similarity with exonic portions of other lncRNAs. To our knowledge, this represents the first draft of a genome-wide catalog of shared lncRNA exons.

### Nucleotide variation in modules

To further investigate whether exon modules may represent conserved functional units, we analyzed the occurrence and frequency of single nucleotide polymorphisms (SNPs) in these regions, as a lower incidence of variants may indicate the existence of constraints associated with functional sequences, due to the effects of purifying selection [34]. Accordingly, we collected SNP data from the 1000 Genome project from dbSNP 153 [35] and we observed 12.87 variants per thousand bases in control exons (which are not modules) and 11.83 in modules. We then obtained from the ALFA allele frequencies aggregator [36] a total of 764,005 SNPs located in lncRNA exons, [36] and their associated frequencies. For each exon, we calculated the index of nucleotide diversity *θπ* [37] as

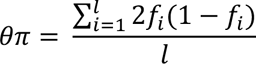

where *fi* represents the frequency of variants in the *i* th position of the exon sequence in the population, and *l* represents the length of the exon.

After comparing the distributions of *θπ* scores with the Mann-Whitney U test, we obtained a p-value of 2.14e-02 in the comparison between modules and exons from functionally annotated genes, a p-value of 2.77e-02 from the comparison between exon modules and other lncRNA exons and a non-significant p-value (7.45e-01) from the comparison between functionally annotated and others, confirming a significant lower propensity to harbor variation in exon modules as compared to the other two groups. These findings indicate the existence of evolutionary constraints which limit the occurrence of variants with polymorphic frequencies in exon modules, which in turn may reduce the rate of evolutionary change in the long-term. We also looked at the frequency of polymorphic complete exon deletions, but the results were not statistically significant (data not shown).

### Search for characteristics shared with protein coding genes

To confirm that exon modules do not simply represent mis-annotated protein domains, we compared their sequence characteristics with those of known coding genes.

França et al. [38] observed that symmetric shuffling units (exons whose length is an exact multiple of three) are strongly over-represented in human protein coding genes, due to their lower impact on the reading frame when transposed. We found an opposite trend in lncRNA exon modules, with only 25% having a length that is a multiple of three, which confirms the lack of relevance of the reading frame. By contrast, in the remainder of the exons, this proportion is 33%, i.e. what would be expected under a random model.

The transition/transversion ratio (Ti/tv) among polymorphic variants should be 0.5 under a purely random model, resulting from four possible transitions/eight possible transversions. However, real data depart remarkably from this expectation, with functional regions and protein coding regions presenting values higher than 0.5, since transitions are more likely to result in non-synonymous substitutions (e.g. when they occur in the third base of a codon) [39]. Exon modules displayed values of 1.9, in line with previous results for lncRNAs [40]. As a reference, these values contrast sharply with those for protein coding genes, which range between 2.8-2.9 +/-0.1 [40].

### Functional hypothesis and organization of putative modules in clusters of lncRNA genes

To further describe exon modules, here we show some examples of their organization within the structure of their lncRNA genes. Only 12 of 218 genes containing exon modules are associated with a known biological function in the lnc2Cancer database [5]. For most of them, the specific region of the lncRNA molecule responsible for that function is unknown. In the next two paragraphs we will provide a more detailed description for two of the identified modules, in an attempt to capture their putative functions. The first example refers to an exon module recognized by virtue of sequence similarity, and the second one refers to an exon module recognized by virtue of structure similarity.

### Identification of a putative YBX1 binding module

Figure 7A (7A Fig.) shows an example of a putative module represented in a pair of exons as a sequence of ∼200 nucleotides sharing a high sequence similarity (>87%). The exons involved are ENSE00003710224.1 and ENSE00003838358.1 which belong to genes ENSG00000182165.17 (also known as TP53TG1) and ENSG00000285540.1, respectively. TP53TG1 is a lncRNA involved in the p53 network response to DNA damage [41], which has a role as tumor suppressor by blocking the tumorigenic activity of the RNA binding protein (RBP) YBX1 [42]. More in detail, the expression of TP53TG1 is induced by p53 under cellular stress conditions that involve the induction of double-strand breaks [41], while the interaction in the cytoplasm between TP53TG1 and YBX1 prevents the migration of the latter inside the nucleus where it might promote the transcription of a series of oncogenes [43]. *Diaz-Lagares et al.* [42] demonstrated that a central region of TP53TG1, which includes the putative module in the exon ENSE00003710224.1, is responsible for YBX1 binding. Moreover, they proved that YBX1 binding motifs CACC are necessary to ensure the tumor-suppressor function of TP53TG1. We identified two occurrences of the CACC motif in ENSE00003710224.1 and one in ENSE00003838358.1, suggesting a common role for this module.

**Fig.7.**
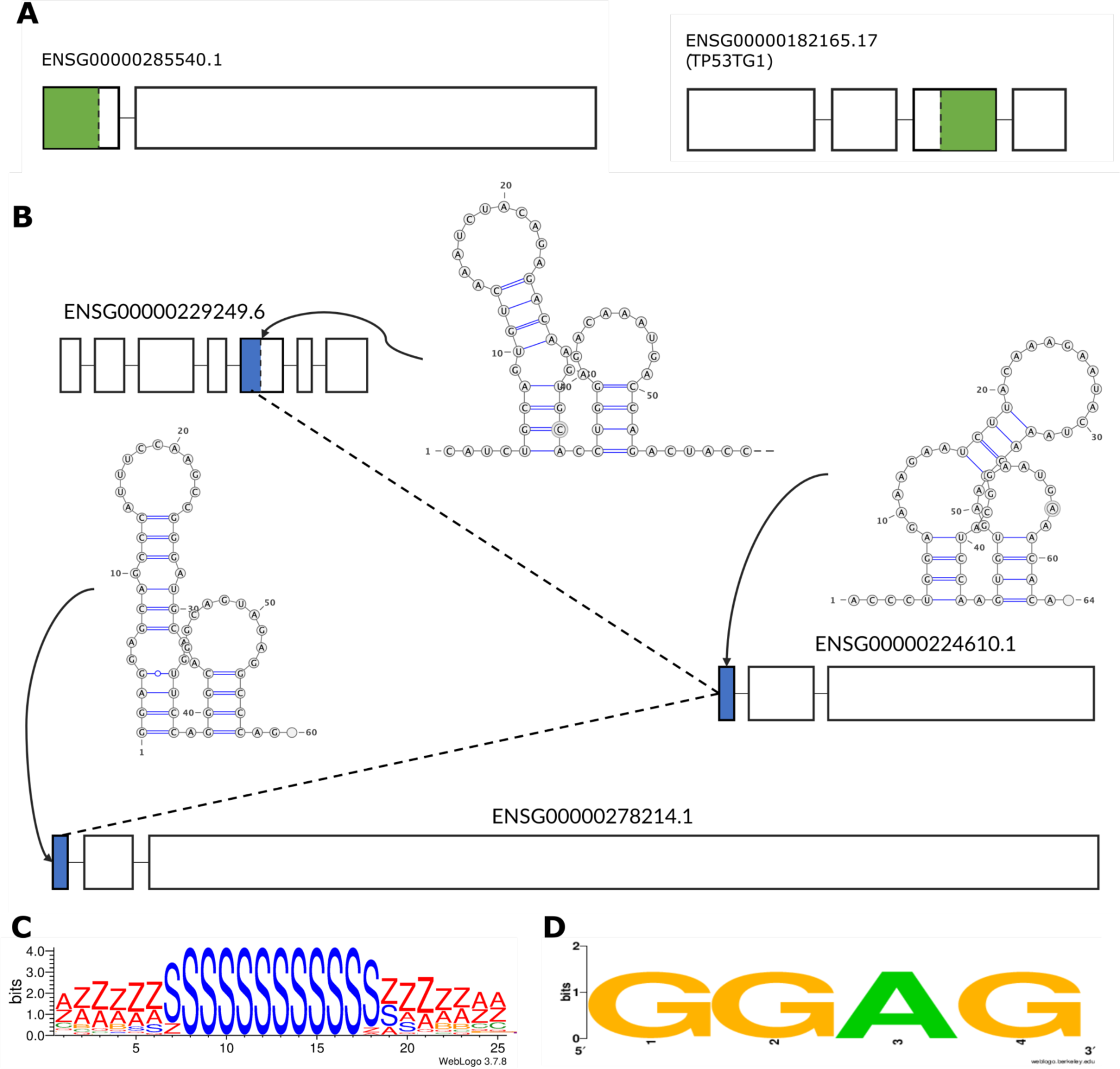
Organization of a sequence and a structure module and identified motifs. A) Schematic representation of the lncRNA genes containing the putative YBX1 binding module (in green); B) Representation of the lncRNA genes containing the exons with the putative LIN28B binding module and their secondary structures. The blue boxes represent the exons with high structural similarity that form the module; C) secondary structure motif revealed by BRIO represented with the BEAR alphabet [50]; D) sequence motif recognized by ZnK in the three modules. The RNA secondary structure representations were generated using VARNA (Darty et al. 2009); Sequence and structure logos were generated using WebLogo [51].

### Identification of a putative LIN28B binding module

Figure 7B-D (7B-D Fig.) shows an example of a module with high structure similarity, embedded in dissimilar sequence contexts. The exons involved are ENSE00003741285.1, ENSE00001800736.1 and ENSE00001782399.1 which belong to ENSG00000278214.1, ENSG00000224610.1 and ENSG00000229249.6, respectively (7B Fig.). These three exons fold into a similar secondary structure, composed of two stems ending with a hairpin loop, with one of the two stems having one or two internal loops.

To detect a possible function, common to the three representatives of this exon module, we searched for the presence of enriched structure and sequence motifs using the BRIO web server (see Materials and Methods). BRIO identified a significantly enriched (Fisher’s exact test padj<0.05) structure motif shared between all the exons of the group (7C Fig.). This particular motif was associated by Adinolfi *et al.* [44] with a series of different RNAs capable of binding some RBPs including LIN28B. This is an evolutionary conserved RBP involved in several cellular processes, which acts as a critical oncogene activated in cancer [45]. LIN28B is known to be able to bind different mRNAs, including a set of mRNAs for splicing factors [46], miRNAs [47] and lncRNAs such as NEAT1 [48]. Furthermore, LIN28B C-terminal zinc knuckle (ZnK) mediates specific binding to a conserved GGAG motif [49] which is also a sequence motif present in all the three representatives of this module (7D Fig.). These observations suggest a possible role of this module in binding LIN28B.

## Discussion

This work identified a set of lncRNA exons with high sequence and/or structure similarity that are embedded within globally dissimilar genes, confirming the hypothesis of exon sharing between this class of molecules similar to protein-coding genes. This set contains a total of 340 pairs of exons that can be grouped, on the basis of their reciprocal connections, in 106 clusters. In contrast to previous work [18], our analysis focused on exons that do not contain repetitive sequences. The resulting dataset of exon modules likely represents the result of recent segmental duplications that are almost exclusively found in humans and higher primates. These findings support the hypothesis that the non-coding transcriptome is structured into modular domains, similar to the organization observed in protein-coding genes.

Approximately 4% (218 out of 5,423) of all the lncRNA genes in our dataset contain an exon module. Even though we cannot assign a specific function to each of these modules, as it has been done for the majority of protein coding domains, it is tempting to infer that sharing of functional modules between different lncRNAs may contribute to expanding the functional repertoire of the non-coding genome, similar to the shuffling of functional exons in coding sequences [12].

LncRNA exon modules identified in this work display a higher degree of sequence conservation and synteny in four primate great ape species than the remainder of lncRNA exons. A high level of conservation between related species is suggestive of purifying selection and is a landmark characteristic of functional genetic elements [52]. Exon modules also harbor a lower frequency of SNPs compared with control sequences, which suggests that purifying selection also persists intra-specifically in human populations. Our set included 46 exon pairs highly similar in both sequence and structure (1F Fig.), which are associated with the highest conservation scores. Even though we cannot infer the age of the duplication/shuffling event based on our analysis, our results show that the exons involved are subjected to extreme purifying selection, which preserved both sequence and structure. Taken together, this evidence suggests that these modules play an important role within their respective lncRNA genes, even though their exact function is yet to be characterized.

Some of the modules may also have undergone an accelerated divergence. Our set includes 219 and 75 exon pairs similar only in sequence or structure, respectively, and since our inclusion criteria considered both similarity and evolutionary conservation, examples of accelerated evolution may have escaped our search. This mechanism is equally relevant, especially when searching for evolutionary innovations specific to the human lineage. However, different methods than those used here are required to identify such cases. Finally, it was reported that homologous lncRNAs can, in some cases, conserve their function over long evolutionary times, despite having diverged in both their nucleotide sequences and their secondary structures [53]. The above considerations suggest that our analysis may underestimate the extent of module sharing in lncRNAs. Other limitations include the fact that the correct identification of exons within lncRNAs is strongly dependent on the reliability of the reconstruction of the whole transcript structure. This is usually summarized by the TSL parameter (Transcript Support Level) which we included, for every exon, in the Supporting information (S1 Table).

In the few cases for which functional information on a lncRNA is available, it may be possible to infer the function of the shared module. We report two examples of modules conserved in either sequence or structure. In both cases, the ability to bind specific targets is the inferred associated function.

Overall, our results highlight the presence of groups of exons sharing high sequence or structure similarity within dissimilar lncRNA genes. These exons are highly conserved across primate species and depleted of inter-individual variation among humans (SNPs), and we suggest that they may represent functional modules.

The identification of these modules could constitute a tool for decoding the function of the many lncRNAs that are currently uncharacterized. Membership in a shared exon cluster represents a feature that deserves annotation, even though conclusive proof of shared function will require experimental evidence.

## Materials and Methods

### Dataset

We used gencode version 29 [3], to select 34,509 exons annotated as long intergenic non-coding RNA, which do not have overlaps with protein coding genes, and downloaded their chromosomal coordinates as a gtf file. We then used these coordinates to obtain the corresponding sequences from the hg38 version of the human genome (UCSC genome browser), converting the gtf to bed file and using the getfasta tool from the bedtools suite [54], with repetitive sequences masked by RepeatMasker (Smit et al., unpublished data, www.repeatmasker.org) and Tandem Repeats Finder [55]. We removed 18,703 exons containing repetitive sequences and retained 15,806 exons. 3,709 of these were shared by different isoforms of the same lncRNA gene. In such cases we only considered the longest isoform, thus obtaining a final set of 12,097 non-overlapping exons that do not contain repetitive sequences. These exons belong to 5,423 different lncRNA genes.

### Sequence alignments

All exon sequences were compared to each other using the Needleman and Wunsch global alignment algorithm [56], using the same default gap penalties scores as the EMBOSS Needle tool for global alignments of nucleic acids sequences [57] (−10 for gap insertions, −0.5 for gap extensions) and the EDNAFULL substitution matrix.

### Structure alignments

The secondary structure of each exon was calculated using RNAfold [57,58], as the minimum free energy (MFE) structure, and represented by its dot-bracket notation. These representations were converted into the BEAR alphabet for RNA secondary structure notation (Mattei et al. 2014). The BEAR alphabet is an encoding method for RNA secondary structure, whose characters encode for a specific secondary structure element (loop, stem, bulge and internal loop) with specific length (e.g. a nucleotide that is part of a stem of length 5 is represented by one character and a different character is used to represent a stem of a different length). The global structure alignments were performed using the BEAGLE algorithm [59], with default parameters (−2 for gap insertions, −0.7 for gap extensions, +0.6 for the sequence match bonus) and the substitution matrix for RNA structural elements (MBR, Matrix of Bear-encoded RNAs) described in [50]. To avoid favoring alignments between unstructured regions we modified the original MBR, assigning a score of 0 to matches in these regions. BEAGLE is an algorithm for pairwise RNA secondary structure global comparison similar to the Needleman and Wunsch algorithm for sequence alignments.

For both sequence and structure alignments we considered the scores of the aligned sequences after trimming external gaps. The score of each alignment was normalized by its length, to avoid biases towards longer sequences. We selected only alignments of a length of at least 50 nucleotides after the external gap trimming. The final distributions consisted of approx. 73 million values, with z-scores ranging from ∼-36 to ∼16 and from ∼-3 to ∼9, respectively.

### Repetitive elements and cis-regulatory elements

Repetitive sequences were mapped using the rmsk table from the UCSC genome browser, which is derived from RepeatMasker (Smit et al., unpublished data, www.repeatmasker.org).

Cis-regulatory elements coordinates are derived from the ENCODE Registry of candidate cis-Regulatory Elements (cCREs) combined from all human cell types [60]. The enrichments are calculated using a Fisher’s exact test between modules containing a particular CRE and the other lncRNA exons of the dataset with a Benjamini-Hochberg correction.

### Evolutionary conservation score

The evolutionary conservation score for each exon was calculated using an approach similar to [61], using the BLAST+ suite of command-line tools [62]. More specifically, the BLASTn algorithm was used to perform an alignment of all the lncRNA exons of our dataset (12,097). In view of the pattern of the evolutionary conservation of lncRNA sequences [14], we used the genomes of four primate species closely related to *H. sapiens*: *Pan troglodytes* (Chimpanzee, taxid:9598), *Pan paniscus* (Bonobo, taxid:9597), *Pongo pygmaeus* (Orangutan, taxid:9601) and *Gorilla gorilla* (Gorilla, taxid:9592). For each lncRNA exon we then calculated a comprehensive conservation score as the sum of the best match bit-score over the four species, divided by the length of the query sequence. Though the four organisms are phyletically correlated, we used this procedure to buffer lineage-specific effects and potential genome annotation errors.

For both sequence and structure similarity scores, the resulting distributions were compared with the inter-specific degree of sequence conservation, under the hypothesis that constraints on exon variation acted both intra- and inter-specifically. These comparisons were used to explore the relationship between intra- and inter-specific conservation scores around the z-score value of 6.0 proposed by [63] as the threshold to distinguish homologous sequences (1 Fig.).

We excluded from this comparison exon pairs located in genes that are globally similar as the similarity of the exons would simply reflect gene paralogy. To do so we performed a pairwise alignment of the genes containing the exon pairs using BLASTn. The genomic coordinates of the whole genes, including the introns, were retrieved from the gencode version 29 gtf file [3], and we used the same procedure described above for the exons to obtain their sequences. Local alignments were performed considering the smallest gene of the pair as the query and the longest as the subject, and excluding pairs presenting a total query coverage greater than or equal to 80%. For each exon pair, we also checked the coordinates from the bed file, excluding overlapping pairs.

### Syntenies

Synteny data were collected from SyntDB [32], which takes into account positional conservation and sequence similarity to identify syntenic regions of human lncRNAs across primates. This database comprises synteny information for 55632 transcripts. From this dataset we selected conservation data in Chimpanzee, Bonobo, Orangutan and Gorilla for the 8,390 lncRNA transcripts containing the 12,097 exons in our dataset.

### Single nucleotide polymorphisms (SNPs)

SNPs locations were retrieved from common dbSNP 153 (variants with a minor allele frequency (MAF) of at least 1% (0.01) in the 1000 Genomes Phase 3 dataset) [35] and population frequencies were obtained from the ALFA allele frequency aggregator project [36]. The release 2 vcf format file contains variant frequency data aggregated from 79 different studies on more than 900 million SNPs. We used the tabix tool from the SAMtools suite of programs [64] to select SNPs located within one of the 12,097 exons in our dataset, obtaining ∼764,000 variants with associated allele frequency information.

### Transition/transversion ratio

The transitions to transversion ratio (Ti/Tv) was calculated by using the variant data present in the common dbSNP 153 (see above) for all the 12,097 lncRNA exons in our dataset, as the number of pyrimidine-pyrimidine or purine-purine substitutions (transitions), divided by the number of purine-pyrimidine or pyrimidine-purine substitutions (transversions).

### Protein coding exons

The protein coding exon coordinates were obtained from the gencode version 29 annotation and mapped on the hg38 version of the human genome using the same procedure described for the lncRNA exons.

### Motifs scan

The search for sequence and structure motifs in the putative LIN28B binding module was performed using the BRIO (BEAM RNA Interaction mOtifs) web server [65]. This tool enables the identification of RNA sequence and structure motifs involved in protein binding in one or more input RNA molecules, by measuring, through a Fisher’s exact test, their enrichment compared to a background of RNAs from Rfam with similar length and structure content, defined as the fraction of paired nucleotides in the RNA secondary structure. The database of motifs that is included in BRIO is derived from high throughput protein-RNA binding experiments (PAR-CLIP, eCLIP and HITS) analyzed in [44]. For this analysis, we considered the default enrichment significance threshold of p-value<0.05 to evaluate the enrichment of a motif in a group of exon modules. We chose to use this algorithm because in addition to identifying common motifs on some particular modules, it allows us to associate them with motifs enriched in RNA that interact with specific proteins from experimental data.

## Author Contributions

Conceptualization: Francesco Ballesio, Gerardo Pepe, Gabriele Ausiello, Andrea Novelletto, Manuela Helmer-Citterich, Pier Federico Gherardini.

Data curation: Francesco Ballesio.

Investigation: Francesco Ballesio.

Funding acquisition: Manuela Helmer-Citterich, Pier Federico Gherardini.

Writing – original draft: Francesco Ballesio.

Writing – review & editing: Gerardo Pepe, Gabriele Ausiello, Manuela Helmer-Citterich, Pier Federico Gherardini, Andrea Novelletto.

Gabriele Ausiello, Manuela Helmer-Citterich, Pier Federico Gherardini, Andrea Novelletto are senior authors.

## Data Availability

The annotation was obtained from GENCODE v29 (https://www.gencodegenes.org/human/release_29.html).

The hg38 version of the human genome was downloaded from UCSC genome browser (http://hgdownload.cse.ucsc.edu/goldenPath/hg38/bigZips/).

The BEAGLE webserver for RNA structure alignments is available at: http://beagle.bio.uniroma2.it.

Functionally annotated lncRNA was downloaded from: http://bio-bigdata.hrbmu.edu.cn/lnc2cancer.

Variant frequencies in human populations are available in the ncbi website (https://www.ncbi.nlm.nih.gov/snp/docs/gsr/alfa/#ftp-download).

The BRIO webserver for RNA interaction motif search is available at: http://brio.bio.uniroma2.it.

SynthDB is available at: http://syntdb.amu.edu.pl.

For a list of the 340 exon pairs identified see S1 Table.

## Supporting information

S1 Table A list of identified exonic modules and their properties.

S2 Table Repetitive elements in the regions flanking the exonic modules.

S3 Table Percentage of transposase domains identified in genes containing exon modules and in other lncRNA genes.

---

**Fig. S1.**
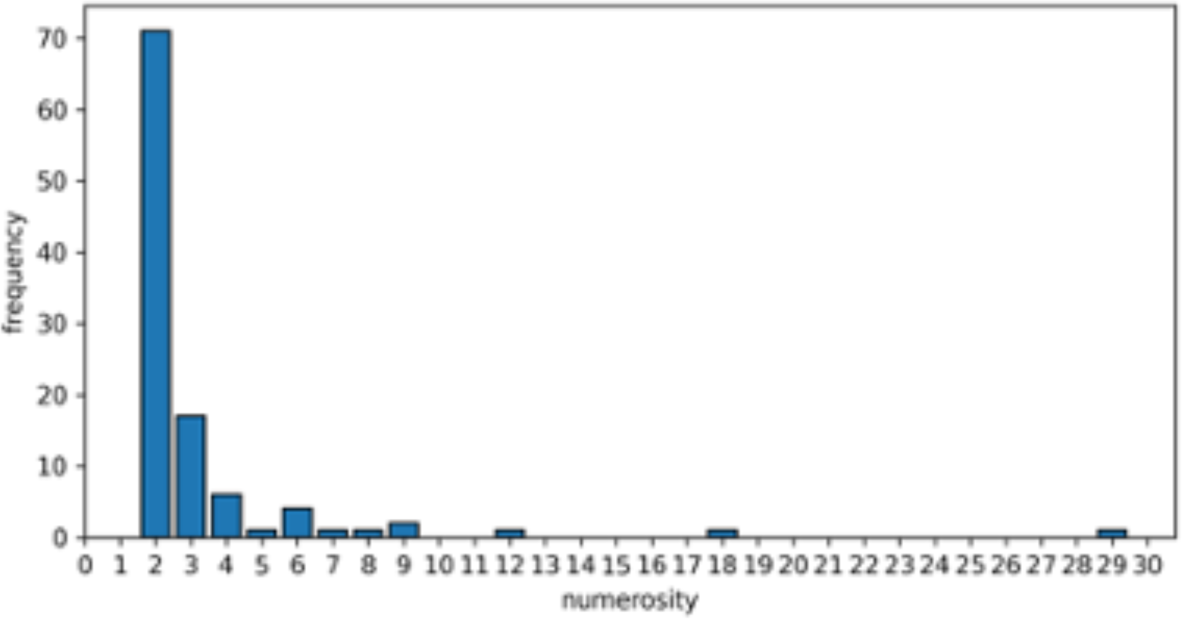
Numerosity of lncRNA exons per exon cluster.

**Fig. S2.**
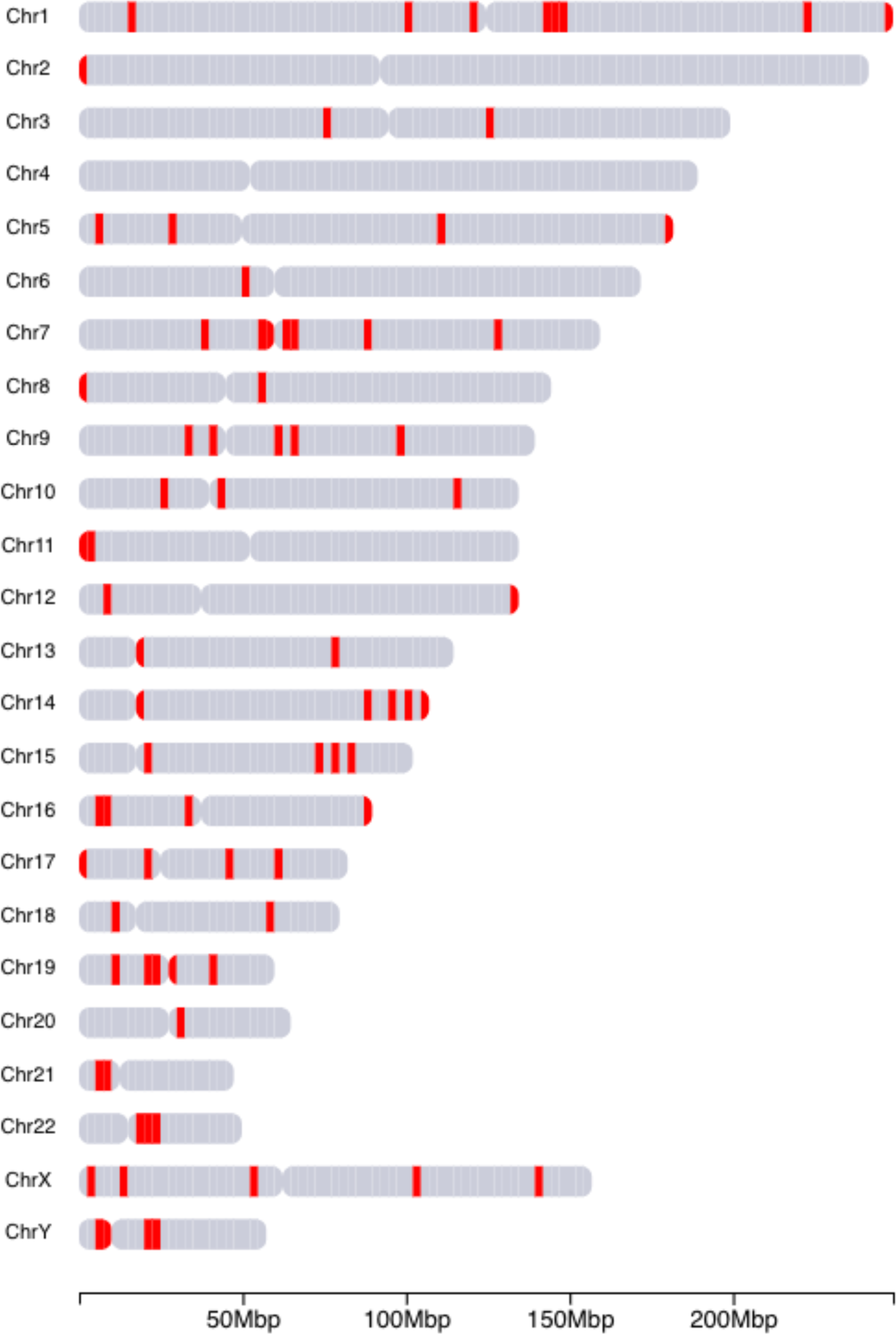
Positions of the exon modules on the human chromosomes

**Fig. S3.**
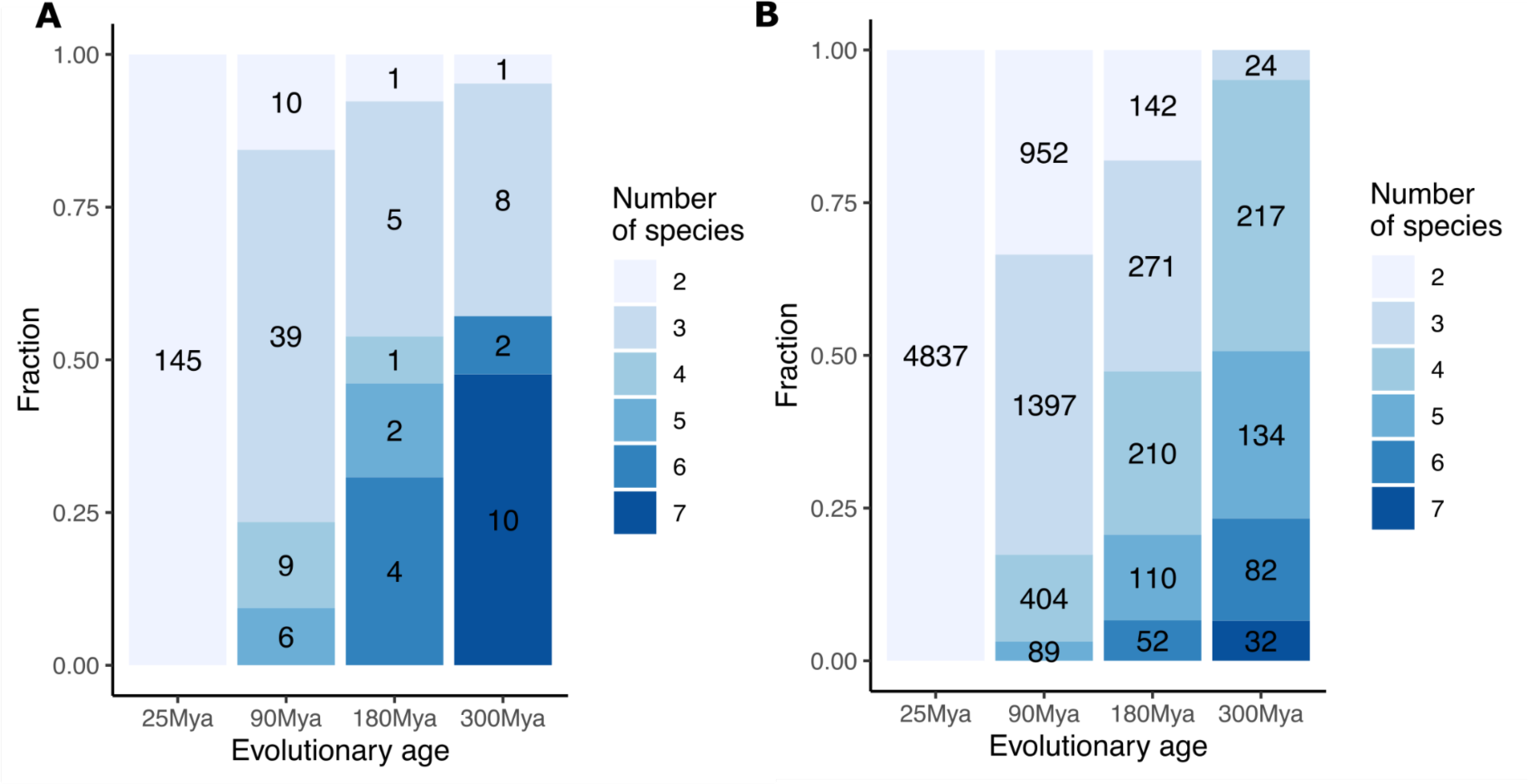
Comparison of BLAST hit frequencies at different evolutionary divergence ages of exonic modules (A) and lncRNAs genes analyzed by Sarropoulos et al. (B).

